# Kinetic modelling of sterol transport between plasma membrane and endo-lysosomes based on quantitative fluorescence and X-ray imaging data

**DOI:** 10.1101/2023.01.28.526075

**Authors:** Daniel Wüstner, Alice Dupont Juhl, Jacob Marcus Egebjerg, Stephan Werner, James McNally, Gerd Schneider

## Abstract

Niemann Pick type C1 and C2 (NPC1 and NPC2) are two sterol-binding proteins which, together, orchestrate cholesterol transport through late endosomes and lysosomes (LE/LYSs). NPC2 can facilitate sterol exchange between model membranes severalfold, but how this is connected to its function in cells is poorly understood. Using fluorescent analogs of cholesterol and quantitative fluorescence microscopy, we have recently measured the transport kinetics of sterol between plasma membrane (PM), recycling endosomes (REs) and LE/LYSs in control and NPC2 deficient fibroblasts. Here, we employ kinetic modeling of this data to determine rate constants for sterol transport between intracellular compartments. Our model predicts that sterol is trapped in intraluminal vesicles (ILVs) of LE/LYSs in the absence of NPC2, causing delayed sterol export from LE/LYSs in NPC2 deficient fibroblasts. Using soft X-ray tomography, we confirm, that LE/LYSs of NPC2 deficient cells but not of control cells contain enlarged, carbon-rich intraluminal vesicular structures, supporting our model prediction of lipid accumulation in ILVs. By including sterol export via exocytosis of ILVs as exosomes and by release of vesicles – ectosomes – from the PM, we can reconcile measured sterol efflux kinetics and show that both pathways can be reciprocally regulated by the intraluminal sterol transfer activity of NPC2 inside LE/LYSs. Our results thereby connect the in vitro function of NPC2 as sterol transfer protein between membranes with its in vivo function.

## 1. Introduction

Late endosomes and lysosomes (LE/LYSs) play important roles in degradation of biomolecules and in sensing the nutritional status of the cell (Appelqvist et al., 2013;Settembre et al., 2013). To exert such functions, the abundance of cholesterol and other lipids in the limiting membrane as well as in internal vesicles of LE/LYSs is crucial (Chevallier et al., 2008). Lysosomal cholesterol accumulation impairs membrane fusion and luminal acidification, attenuates cellular death pathways and affects virus entry and replication (Cox et al., 2007;Fraldi et al., 2010;Appelqvist et al., 2011;Carette et al., 2011;Stoeck et al., 2018). Mammalian cells receive most of their cholesterol from receptor-mediated endocytosis of low density lipoprotein (LDL), which dissociates from its receptor in early endosomes followed by trafficking of LDL to LE/LYSs for degradation (Goldstein and Brown, 2009;Luo et al., 2020). Regulation of this trafficking pathway is complex and ensures sufficient cholesterol supply but also prevention of cholesterol accumulation (Goldstein et al., 2006;Juhl and Wüstner, 2022).

Several proteins have been involved in export of endogenous and LDL derived cholesterol from LE/LYSs (Luo et al., 2020;Juhl and Wüstner, 2022;Lu, 2022). Among them, Niemann Pick C1 and C2 (NPC1 and NPC2) play a central role, and mutations in either protein can lead to pronounced accumulation of cholesterol and other lipids in LE/LYSs of various organs, including the brain – a hallmark of Niemann Pick type C disease. NPC1 is a large transmembrane protein in the limiting membrane of endo-lysosomes with at least two sterol binding sites; one in the luminal N-terminal domain (NTD) and the other in the transmembrane sterol sensing domain (SSD), respectively (Pfeffer, 2019). NPC2 is a small sterol-binding protein in the lumen of LE/LYSs which has been shown to facilitate sterol exchange between model membranes (Xu et al., 2008;McCauliff et al., 2011). Mammalian and yeast NPC2 can transfer sterol to the NTD of NPC1 (named NCR1 in yeast), suggesting a mechanism for sterol transfer between both proteins in the LE/LYSs (Infante et al., 2008;Wang et al., 2010;Winkler et al., 2019). Once bound to the NTD of NPC1 (or NCR1), cholesterol is supposed to move through a tunnel in the protein to get finally inserted into the lysosomal membrane. Such a tunnel was first identified in the yeast homologue NCR1 and later also in mammalian NPC1, and key residues important for sterol transport could be identified (Winkler et al., 2019;Long et al., 2020;Qian et al., 2020;Saha et al., 2020). NPC2 has been shown to bind to NPC1 but also to other abundant proteins in endo-lysosomes, such as LAMP-2 (Deffieu and Pfeffer, 2011;Li and Pfeffer, 2016;Li et al., 2016). Thus, NPC2 might deliver cholesterol and other sterols not only to NPC1 but also to LAMP-2 and other potential sterol transporters in the lysosomal membrane. Apart from ergosterol, the yeast NPC2 binds a variety of other lipids including analogues of phosphatidylserine, phosphatidylcholine, and phosphatidylinositol, while mammalian NPC2 binds lysobisphosphatidic acid (LBPA), a lipid specifically found in intraluminal vesicles (ILVs) inside LE/LYSs (McCauliff et al., 2019;Moesgaard et al., 2020). LBPA enhances sterol exchange between vesicles by NPC2 in vitro (Cheruku et al., 2006;McCauliff et al., 2015). Adding LBPA or its metabolic precursor phosphatiyl glycerol (PG) to fibroblasts facilitates cholesterol export from LE/LYS, even in NPC1-deficient fibroblasts (Ilnytska et al., 2021a). Thus, NPC2 could be a general lipid carrier inside endo-lysosomes, and its function seems to depend on other lipids, such as LBPA. How NPC2’s known ability to facilitate sterol transfer between liposomes relates to its in vivo function in LE/LYSs is not fully understood. We have recently shown that sterol trafficking from the PM to LE/LYSs is altered in NPC2 deficient fibroblasts, and that lack of NPC2 results in endo-lysosomal trapping of the fluorescent cholesterol analogue dehydroergosterol (DHE) (Berzina et al., 2018). DHE is intrinsically fluorescent with no dye molecule attached to it and therefore resembles cholesterol very closely. Internalization of purified NPC2 can mobilize this sterol pool and causes reallocation of LE/LYSs to the PM from where the excess cholesterol can be released (Juhl et al., 2021b). We also showed that this last step is facilitated by ABCA1/Apoprotein A1 and involves shedding of sterol-rich vesicles from the PM (Juhl et al., 2021b).

While we could demonstrate that internalized NPC2 co-localizes extensively with fluorescent cholesterol analogues during efflux, we do not know, exactly how it could activate the trapped sterol pool inside LE/LYSs. To answer this question, we combine here mathematical modeling of the previously published sterol transport data with X-ray microscopy of the ultrastructure of endo-lysosomes of control and NPC2-deficient cells. Our modelling analysis and experimental results can explain the observed defects in intracellular sterol transport and efflux in NPC2-deficient cells as a consequence of impaired sterol transfer from ILVs to the limiting membrane of LE/LYSs. This leads to an expansion of the sterol pool in ILVs which can be released from cells as exosomes by lysosomal exocytosis. Our model analysis shows that this sterol efflux pathway could dominate in NPC2-deficient cells, while in control cells, the more efficient delivery of sterol to the PM allows for preferred sterol efflux from the cell surface. Efflux of cholesterol from the PM could be mediated by release of vesicles, so-called ectosomes, as well as by extracellular acceptor proteins, like apoprotein A1 (apoA1) and albumin. Our study provides an integrated model combining the established in-vitro function of NPC2 as sterol transfer protein and lipid solubilizer with its in vivo function in regulating cholesterol flux from endo-lysosomes.

## 2. Methods

### 2.1. Reagents, cell culture and labeling for X-ray microscopy

Human skin fibroblasts from control subject (Coriell Institute #GM08680) were from a male healthy donor, while NPC2 deficient human skin fibroblasts were from Coriell Institute #GM18455 (a male patient affected by two point mutations at the NPC2 locus resulting in a nonsense mutation at codon 20 in allele 1 and in a missense mutation at codon 47 of allele 2, respectively). They were grown at 37 ◦C in an atmosphere of 5% CO2 until 90 % confluence in complete DMEM culture medium (from GIBCO BRL; Life Technologies, Paisley, Scotland) supplemented with 1% glutamine, 1% penicillin and 20 % FBS for diseased cells or 10 % FBS for control cells. TopFluor-cholesterol (TF-Chol) and other reagents were from SIGMA (Denmark). Buffer medium contained 150 mM NaCl, 5 mM KCl, 1 mM CaCl2, 1 mM MgCl2, 5 mM glucose and 20 mM HEPES (pH 7.4). TF-Chol was loaded onto methyl-β-Cyclodextrin to form a complex as described (Juhl et al., 2021b). Briefly, 1.5 μM of TF-Chol from an ethanol stock were dried under nitrogen and mixed with phosphate buffer solution (PBS) containing 1 mg BSA and 40 mg MCD to a final volume of 2 ml under rigorous vortexing for 5 min. Cells were washed with buffer medium before they were pulse labeled for 3 min with 100 μL TF-Chol/MCD solution, washed and chased for 2h at 37 ◦C, in buffer medium.

### 2.2. Generation of fluorescence time courses as input for modeling, data regression and calculations

Fractional fluorescence of the cholesterol analog DHE in the PM, REs and LE/LYSs was measured in fibroblasts from healthy subjects or from NPC2 disease patients, as described in our previous publication (Berzina et al., 2018). Quantification was based on image segmentation using organelle-specific markers and the ImageJ program with an in-house developed plugin (Rueden et al., 2017;Berzina et al., 2018). For multi-compartment parameter estimation either the SAAM software or the Symfit python module was used (Barrett et al., 1998). Steady states of the dynamic system and the differential equation for the Weibull model were calculated using Mathematica (Wolfram Research Inc., USA) or SymPy, a python library for symbolic calculations (Meurer et al., 2017). Numeric simulations of the ordinary differential equation systems that we derived were implemented either in Mathematica or in Python using the odeint module of SciPy (Virtanen et al., 2020).

### 2.2. Soft X-ray tomography of human fibroblasts

Healthy and NPC2 deficient human fibroblasts were grown to a confluency of 90%, and, after trypsin treatment, split onto Poly-D-Lysine coated R 2/2 grids (QUANTIFOIL 100 Holey Carbon Films, Grids: HZB-2 Au), which were fixed to the bottom of 12 well plates. Cells were allowed to settle for another 48-72 h before further treatment and fixed with 4% PFA. In some experiments, cells were labeled with TF-Chol loaded onto cyclodextrin for 3 min at 37℃ and chased for 2h before fixation and imaging as described (Juhl et al., 2021b). The cells were kept in 1xPBS until cryo-plunge freezing with liquid ethane and subsequently stored and imaged under liquid nitrogen temperature. Prior to the plunge freezing, a small volume of ∼270 nm silica beads with an outer gold shelf were added to the samples, to serve as fiducial markers for tomographic reconstruction. Soft X-ray tomography (SXT) was performed at Bessy II electron storage ring at the Helmholtz Zentrum Berlin. The SXT data was collected at the full-field transmission X-ray microscope at the beamline U41-PGM, with an X-ray energy of 510 eV and a 25 nm zone plate. Cells were kept under liquid nitrogen temperature during imaging and imaged over a range of 120-125° tilt angels with 1° step size. To collect the fluorescent signal, a widefield microscope connected to the X-ray microscope, with an 100x objective, NA =0.7, was used. The image pixel size was 9.8 nm for SXT and 148 nm for fluorescence microscopy. For alignment of tomogram frames, B-soft was used, while the Tomo3D software was used for reconstructions based on the filtered back-projection algorithm (Heymann and Belnap, 2007;Agulleiro and Fernandez, 2015). Sum projections of selected frames along the reconstructed 3D stack were calculated with in-house developed Macro scripts to ImageJ (Macro Language (nih.gov)). Registration of fluorescence and X-ray images was done in Icy using the ec-CLEM plugin with 3-5 image points as reference markers (ec-CLEM | – Open Source Image Processing Software (bioimageanalysis.org)).

## 3. Results

### 3.1. Four-compartment model of sterol transport reconciles defect in NPC2-deficient cells

There have been several attempts to model the transport defects observed in human fibroblasts lacking functional NPC1; Neufeld and co-workers used compartment modeling to predict impaired recycling from a late endosomal compartment proximal to lysosomes back to the PM in NPC1 deficient cells (Neufeld et al., 1999). Lange and Steck compared cholesterol transport in fibroblasts from healthy subjects and NPC1 disease patients using cholesterol isotopes and subcellular fractionation (Lange et al., 1998). They proposed a simple kinetic model to explain their data in which they assumed that cholesterol exchanges bi-directionally between the PM and the LE/LYSs (Lange et al., 1998). This model was based on studies with radioactive cholesterol which provided evidence for recycling of PM-derived sterol from LE/LYSs (Lange et al., 1997;Lange et al., 2000;Lange et al., 2002). They found that the rate constant describing transport from the PM to LE/LYS (𝑘_𝑃𝑀→𝐿𝐸/𝐿𝑌𝑆_) is increased in NPC1 disease compared to control cells, while the transport in the opposite direction (i.e., from the LE/LYS to the PM; 𝑘_𝐿𝐸/𝐿𝑌𝑆→𝑃𝑀_) is slowed down. Together, this resulted in a 2-3-fold expanded sterol pool at a steady state in the NPC1 disease cells with no change in the ER pool size (Lange et al., 1998). In an attempt to this model for NPC2-deficient cells, we used live-cell imaging of the cholesterol analog DHE and found evidence for impaired sterol export from LE/LYSs back to the PM but not for enhanced sterol endocytosis (Berzina et al., 2018). In addition, trafficking of endo-lysosomes was slowed down, and non-vesicular sterol transport reduced in such NPC2-deficient cells (Lund et al., 2014;Berzina et al., 2018). Incubating those cells with purified NPC2 rescued the lysosomal sterol storage phenotype and increased directional transport of LE/LYS towards the PM (Lund et al., 2014;Juhl et al., 2021b). It has been shown in several cell lines that fluorescent close cholesterol analogues, such as DHE are also transported from the PM to early endosomes comprised of sorting endosomes and the endocytic recycling compartment, here abbreviated as recycling endosomes (REs) (Hao et al., 2002;Wüstner et al., 2002;Wüstner et al., 2005;Petersen et al., 2008). REs ensure efficient recycling of endocytic receptors and ligands, such as transferrin and its receptor in two circuits and were shown to contain a lot of cholesterol in several studies (Gagescu et al., 2000;Hölttä-Vuori, 2002;Möbius et al., 2003b;Maxfield and McGraw, 2004;Mesmin et al., 2011). We found that the close cholesterol analogue DHE is rapidly transported from the PM in human fibroblasts, as well, irrespective of whether functional NPC2 is expressed in those cells or not (Berzina et al., 2018). Sterol trafficking to REs has not been included in the model presented previously by Lange and co-workers, which only considered bi-directional sterol exchange between PM and LE/LYS (see Introduction) (Lange et al., 1998).

Here, we attempted to extend the model developed by Lange & Steck to include transport through REs and also to account for the observation, that sterol efflux from cells is often found to be biphasic suggesting some type of compartmental system with lags for sterol efflux (Haynes et al., 2000;Hao et al., 2002). Mathematically, any compartment system with time lags can be adequately described by assuming one or several intermediate compartments (which is sometimes called the ‘linear chain trick’) (Jacquez and Simon, 2002). The mathematical model, we present here, combines observations on lysosomal sterol trafficking from the literature with our published data by considering sterol transport from the PM (with sterol amount *n*_1_) to the REs (with sterol amount *n*_2_) and further to LE/LYSs (with sterol amount *n*_3_) and from there back to the PM (Berzina et al., 2018). Thus, we model also sterol cycling between PM and LE/LYSs but consider REs as an intermediate compartment. We did not include several other compartments in our model because current data indicates they contribute only a small amount to sterol exchange. Specifically, we do not consider the ER sterol pool nor sterol esters, since only very little sterol is transported to acyl-coenzyme A acyl transferase (ACAT) in the ER in our experiments (i.e., about 1.5% of total DHE and ca. 6% and 0.4% of internalized radioactive cholesterol in control and disease cells, respectively, as judged by lipidomics analysis of sterol esterification) (Berzina et al., 2018). While this sterol pool is important for regulation of cholesterol synthesis and expression of sterol transporters (Infante and Radhakrishnan, 2017), it is not relevant in the context of our analysis. Also, the trafficking route of some sterol to the trans-Golgi network (TGN) is not considered in our model, as we did not observe significant enrichment of PM-derived DHE in the TGN in our experiments in human fibroblasts or other cell types (Hao et al., 2002;Wüstner et al., 2002;Wüstner et al., 2005;Berzina et al., 2018). A minor portion of sterol also reaches mitochondria in an NPC2 dependent manner, and we were able to visualize this pool in our recent study using live-cell imaging of cholestatrienol (CTL), another close fluorescent analog of cholesterol (Juhl et al., 2021a). However, the cholesterol content of mitochondria is very low, in part due to its rapid conversion to oxysterols in fibroblasts (Lange et al., 2009). Supporting this notion, we found only faint labeling of mitochondria with CTL derived from the PM, and even though this transport depends on NPC2, it does not contribute significantly to the total sterol flux between the PM and endo-lysosomes (Juhl et al., 2021a). It was therefore not included in our model, which is shown in Fig. 1A with PM and REs as compartment 1 and 2, respectively. All sterol tracer, here assumed to be the fluorescent cholesterol analogue DHE, is initially inserted only in the PM, which is ensured by a short pulse-labeling of human fibroblasts with DHE loaded onto cyclodextrin (Berzina et al., 2018). After washing cells, the transport kinetics of DHE to REs labeled with fluorescent transferrin and of LE/LYSs, stained with fluorescent dextran, is followed over time (Berzina et al., 2018). To describe the observed transport kinetics of DHE in cells lacking functional NPC2, we consider the LE/LYSs as being sub-compartmentalized into compartments 3 and 4, resembling the limiting endo-lysosomal membrane and the ILVs, respectively (Fig. 1A). The need for including a 4^th^ compartment arose from the fact that the observed accumulation of DHE in LE/LYSs consists of two clearly separable phases in the disease cells (see Fig. 1D, green circles). ILVs fulfil diverse functions in endosome maturation, protein degradation and lipid metabolism, and they are the precursor of secreted vesicles called exosomes (Gallala et al., 2011;Huotari and Helenius, 2011;Record et al., 2014). ILVs can be enriched in cholesterol, as shown by immunoelectron microscopy of sterol-binding perifringolysin derivatives (Möbius et al., 2002;Möbius et al., 2003a). Also, in vitro evidence strongly support the hypothesis that NPC2 shuttles sterols between ILVs and the limiting membrane of LE/LYSs either to NPC1 or eventually bypassing NPC1 in the lysosomal membrane (Cheruku et al., 2006;Wang et al., 2010). Based on all these observations, we suggest that NPC2 regulates sterol exchange between compartment 3 (limiting membrane of LE/LYSs) and compartment 4 (ILVs). Absence of NPC2 leads to a biphasic increase of DHE in endo-lysosomes due to accumulation of the sterol in ILVs. Two kinetically defined sterol pools could in principle also relate to separate subpopulations of LE/LYSs (Goldman and Krise, 2010;van der Kant et al., 2013). However, our co-localization studies do not indicate that, as we found DHE enriched in all acidic endosomes in disease fibroblasts (Berzina et al., 2018). Importantly, transport between PM, REs and LE/LYSs is modeled as unidirectional in our model, which simplifies the parametrization significantly. Still, the overall transport between PM and LE/LYS is reversible, accounting for the experimentally observed sterol cycling. The system of ordinary differential equations (ODE) describing the whole model is given by

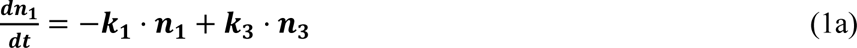

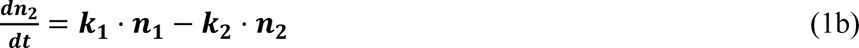

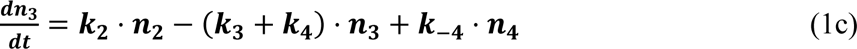

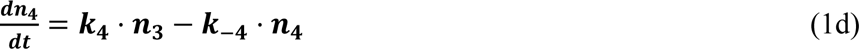

**Figure 1.**
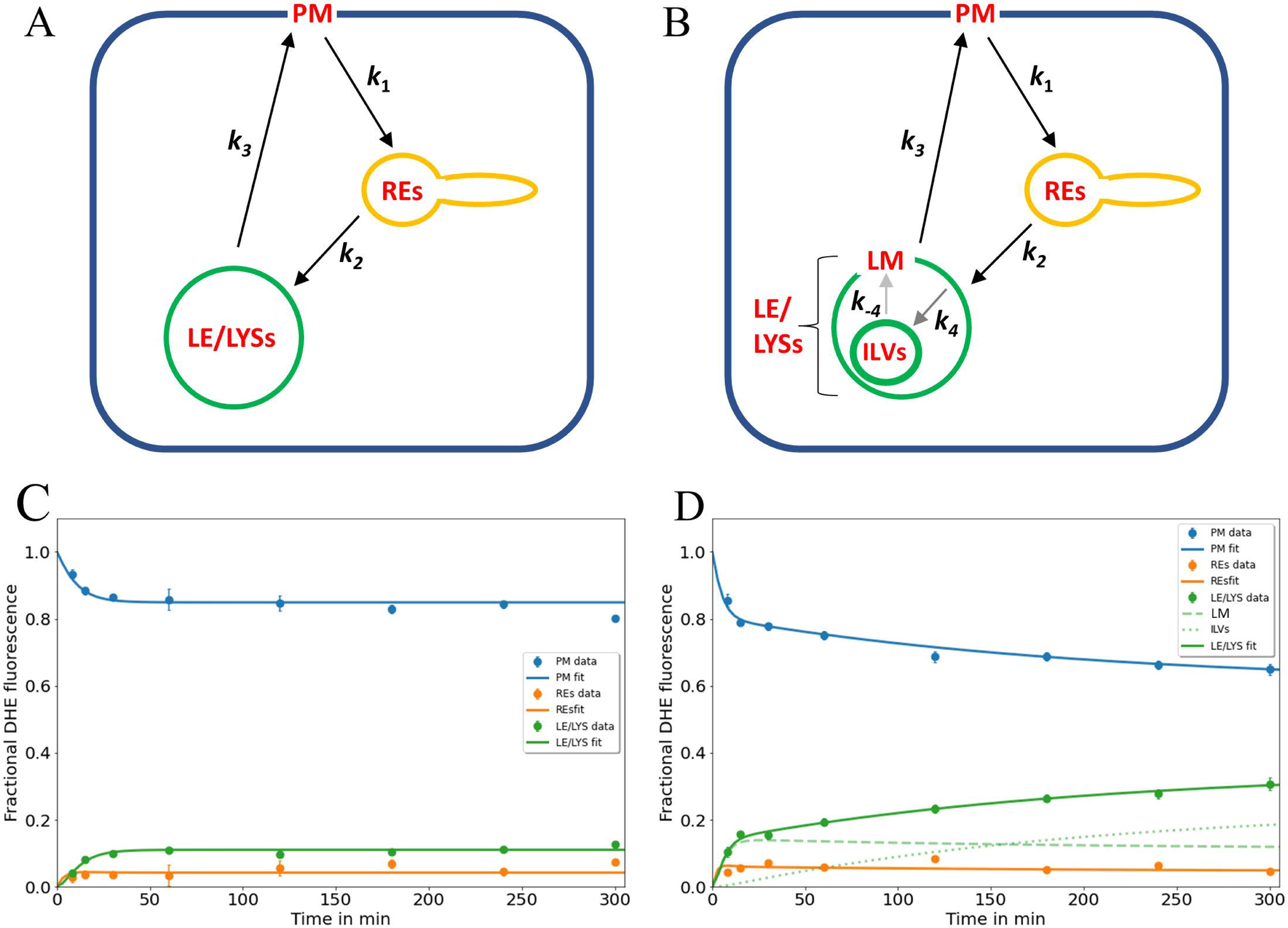
Kinetic modelling of sterol circulation between plasma membrane and LE/LYSs. A, sketch of the model describing sterol transport in control cells with sterol circulation between plasma membrane (PM; compartment 1), the recycling endosomes (REs; compartment 2) and late endosomes/lysosomes (LE/LYSs; compartment 3). The compartments are connected as indicated by the arrows, and sterol transport between them is parametrized by the given rate constants. B, to model the pulse-chase data for disease cells, the LE/LYSs must be subdivided into the lysosomal-membrane (LM, compartment 3) and intraluminal vesicles (ILVs, compartment 4), as otherwise the bi-phasic sterol accumulation in LE/LYSs cannot be explained, B. Sterol exchange between the 3^rd^ and 4^th^ compartment is bi-directional (with rate constants *k*_4_ and *k*_-4_, respectively), but experimentally, we can only assess the sum of both compartments as LE/LYSs. In disease fibroblasts, the bidirectional exchange between limiting membrane and internal vesicles of LE/LYSs is assumed to be very slow due to lack of a functional NPC2 protein (grey arrows between LM and ILVs in B). C, D, fitting of the models to the experimental time courses of DHE transport in control cells (C) and disease cells (D), respectively. The data was generated in our previous publication (Berzina et al., 2018). Results are shown for DHE in the PM (blue symbols, data; blue line, fit), for REs (orange symbols data; orange line, fit) and for LE/LYSs (green symbols, data; green straight line fit). For disease cells, LE/LYSs are modelled as sub-compartmentalized into limiting membrane (LM, compartment 3; green dashed line in D and thin green line in B) and ILVs (green dotted line in D and thick green line in B). See text for further information and Tab. 1 for derived parameters.

The amount of sterol in the *i*^th^ compartment is given by *n*_i_ (with *i*=1, …, 4), and it is assumed that the abundance of DHE in each membrane compartment is proportional to the integrated fluorescence intensity in this compartment. Thus, we directly modelled the fluorescence time courses without conversion into concentration units for which one would need to know the exact compartment volumes. Since those volumes are very difficult to obtain from fluorescence micrographs and because the rate constants, we aimed to infer from our model are first order (i.e., with units of per sec), this option has been neglected. The rate constants describing individual transport steps are defined in Fig. 1A. For modeling the pulse-chase experiment, all sterol was initially in the PM (i.e., *n*_1_(*t*=0) = 1; *n*_2_(*t*=0) = *n*_3_(*t*=0) = *n*_4_(*t*=0) = 0). The ODE system in Eq. 1a-d was solved numerically and the model solutions were fitted to the experimental time courses by a global non-linear regression implemented in SAAM (SAAM Institute, Seattle, WA, USA), as described previously (Wüstner, 2005b). That means parameter sets are determined in parallel from all experimental time courses for control and disease fibroblasts, respectively. In some earlier applications of this approach, we showed that global fitting using SAAM provides high reliability in the fitting performance (Wüstner, 2005b;a;2006b;a). Since we cannot distinguish between the sub-compartments of LE/LYSs in our experiments, the numerical solutions of Eqs 1c and 1d (i.e., for *n*_3_(*t*) and *n*_4_(*t*)) were combined, and this sum was fitted to the time dependent DHE fraction in LE/LYSs. We found that the model shown in Fig. 1A, B accurately describes the experimentally observed time courses including the biphasic behavior of sterol transport from the PM and to the LE/LYSs in the disease cells (Fig. 1D). A compartment model with only three compartments (i.e., *n*_1_(t)-*n*_3_(t), equivalent to cycling of sterol between PM, REs, and the membrane of LE/LYSs) could not account for the dynamics of DHE in disease fibroblasts (not shown) but was sufficient to describe DHE dynamics in control cells (Fig. 1C). A reversible transport scheme, in which sterol shuttles in two independent transport branches to the REs and LE/LYSs, respectively, could also describe the DHE transport data of control cells, but failed to describe the data of disease cells (Fig. S1). Also, using the model of Fig. 1A with only three compartments (equivalent to *k*_4_=*k*_-4_=0) in disease cells provided a less accurate fit in disease cells (not shown), as judged by the Akaike and Bayesian information criteria (Wagenmakers and Farrell, 2004;Daddysman and Fecko, 2013). Including an additional recycling step from REs to the PM did not improve the fit quality but resulted in higher values for the Bayesian information criterion, indicating too many fitting parameters for the available data (not shown). This does not mean, that no sterol recycles from REs to the PM but only that the existing kinetic data in human fibroblasts is insufficient to parametrize this transport step.

Thus, the biphasic sterol accumulation in the LE/LYSs of disease cells is according to our model a consequence of sterol arrival from the REs in the first late-endosomal pool (compartment 3; dashed green line in Fig. 1D) followed by a slow export to the second late-endosomal pool, in which sterol accumulates after some delay (compartment 4; dotted green line in Fig. 1D). We identify the first pool (*n*_3_) with the limiting membrane of LE/LYSs and the second pool (i.e., *n*_4_) with the ILVs. Importantly, transport from the PM to the REs and further to the endo-lysosomal membrane and back to the PM takes place with similar kinetics in control and disease cells (see Tab. 1 for estimated parameter values). Thus, we can allocate the affected transport step in disease cells to the slow cycling of sterol between ILVs and the limiting membrane of LE/LYSs (indicated by grey arrows in Fig. 1B). Given the described function of NPC2 in accelerating sterol exchange between liposomes and shuttling sterol to the limiting membrane of LE/LYSs (Babalola et al., 2007;Infante et al., 2008;Wang et al., 2010;Gallala et al., 2011), our model links the observed in vitro and proposed in vivo function of NPC2.

**Table 1:** Parameters for DHE transport according to the compartment model shown in Fig. 1. Time courses of DHE transport from the PM to REs and LE/LYSs in control and disease cells were fitted to the multi-compartment model shown in Fig. 1 and Eq. 1a-d. Parameters values were optimized in parallel to all compartments by non-linear regression. The mean value, standard deviation (±) and coefficient of variation (CV) of estimated parameters is given, as provided by the SAAM software. For control cells, parameter values for rate constants *k*_4_ and *k*_-4_ could not be adequately fitted, and a three-compartment model was used instead (see text for further details).

| **Rate constant** | $k_1$ (min<sup>-1</sup>)<br>(PM→REs) | $k_2$ (min<sup>-1</sup>)<br>(REs→LE/LYSs) | $k_3$ (min<sup>-1</sup>)<br>(LE/LYSs→PM) | $k_4$ (min<sup>-1</sup>)<br>(LE/LYSs→ILVs) | $k_{-4}$ (min<sup>-1</sup>)<br>(ILVs→LE/LYSs) |
| --- | --- | --- | --- | --- | --- |
| **Control cells** | 0.01204<br>± 0.0012<br>CV=0.0970 | 0.2465<br>± 0.0033<br>CV=0.1355 | 0.09351<br>± 0.011<br>CV=0.1156 | - | - |
| **Disease cells** | 0.03216<br>± 0.0032<br>CV=0.0990 | 0.4331<br>± 0.0049<br>CV=0.1134 | 0.1744<br>± 0.0026<br>CV=0.1495 | 0.0084<br>± 0.0021<br>CV=0.3961 | 0.0039<br>± 0.0015<br>CV=0.2463 |

Previously, we also established a continuous sterol uptake protocol from albumin, in which the sterol donor was continuously present for extended periods of time (Berzina et al., 2018). A particular advantage of the kinetic model shown above is the fact, that it paves the way for a quantitative description of DHE transport using this continuous labeling protocol as well. For that, we have to modify Eq. 1a by adding a constant inflow, *v*_0_, and an efflux rate constant, *k*_5_ (Fig. 2A). For simplicity, we assume sterol efflux taking place from the PM only. Such a pathway is supported by numerous studies and several sterol efflux mechanisms can be envisioned from the PM, such as lipidation of ApoA1 to form high density lipoprotein (HDL) via ABCA1 and other ABC transporters, efflux to albumin, as well as release of sterol-rich vesicles from the PM, so-called ectosomes (Juhl and Wüstner, 2022). We showed recently that the latter pathway contributes to sterol efflux in human fibroblasts, and we demonstrated that upregulation of ABC transporters, such as ABCA1 and co-incubation of cells with ApoA1 enhanced cholesterol efflux from the PM (Juhl et al., 2021b).Thus, NPC2-mediated sterol delivery to the PM and ABCA1/ApoA1could work in tandem to export cholesterol from cells.

**Figure 2.**
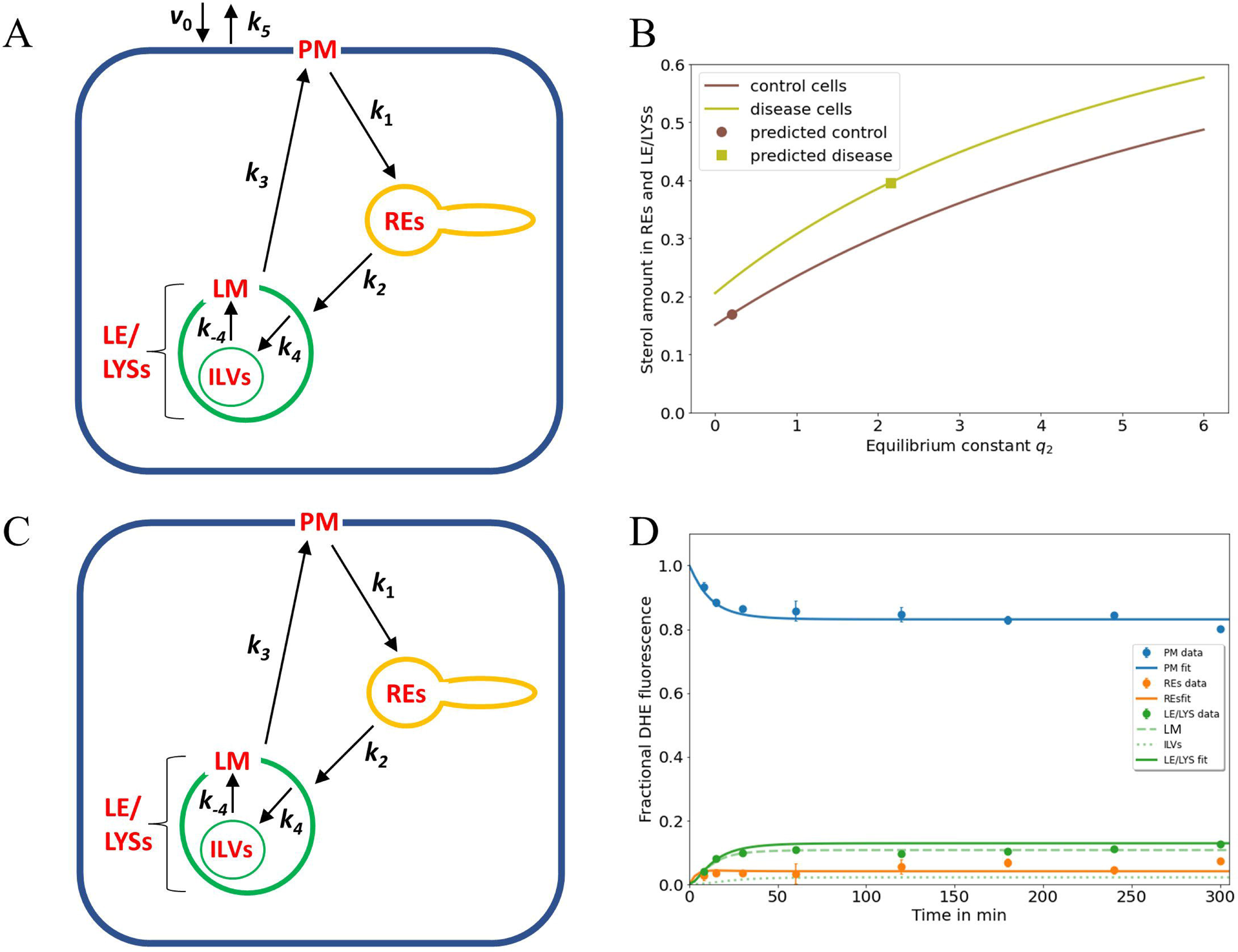
NPC2-mediated sterol transfer between ILVs and limiting membrane of endo-lysosomes determines the extent of intracellular sterol accumulation. A, extension of the kinetic model by including sterol influx with constant rate *v*_0_ and efflux from the PM with rate constant *k*_5_ allows for a steady state analysis (see text and Eqs. 3 to 6). B, the sterol fraction in REs and LE/LYSs relative to total cellular sterol was calculated according to Eq. 6 and is plotted as function of the equilibrium constant between ILVs and the endo-lysosomal membrane (i.e., *q*_2_= *k*_4_ / *k*_-4_) for control cells (brown line) and NPC2-deficient cells (light green line), respectively. The calculated sterol fraction for the value of *q*_2_ obtained by comparing the model with the experiments is indicated by a brown circle for control cells and a light green square for disease cells, respectively. C, D, the full model with sub-compartmentalized LE/LYSs was applied to the data of control cells, assuming that sterol exchange between ILVs (A, C, thin green lines) and the limiting membrane of endo-lysosomes is restored in the presence of NPC2 (A, C, thick green lines, and black arrows inside green LEL/LYSs with rate constants *k*_4_ and *k*_-4_). D, experimental data for intensity of DHE in control fibroblasts (symbols) is compared to the simulated full model with estimated parameter values for rate constants *k*_1_ to *k*_3_ (see Tab. 1) and inferred rate constants *k*_4_ = 0.01686 min^-1^ and *k*_-4_ = 0.0843 min^-1^ corresponding to a ratio of *q*_2_ = *k*_4_ / *k*_-4_ = 0.2 for the sterol fraction. For these conditions, most sterol in LE/LYSs would reside in the limiting membrane (LM, dashed green line) and much less in ILVs (dotted green line) compared to disease cells (see Fig. 1D). See text for further information.

Incorporating sterol uptake and release from the PM into the model leads to a new equation for the PM pool (replacing Eq. 1a) according to (Fig. 2A):

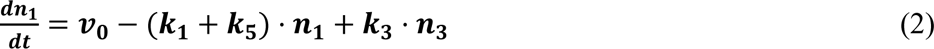

Here, *v*_0_ is the constant rate of sterol uptake, which can be seen as product of a rate constant, *k*_0_, times a constant sterol amount, *n*_0_, bound to albumin in the extracellular reservoir (i.e., *v*_0_ = *k*_0_·*n*_0_). Since we consider this extracellular sterol pool to be in large excess of all cell-associated DHE, the influx *v*_0_ is taken to be a constant. Sterol release from the PM, on the other hand, is proportional to the sterol abundance in the PM with the rate constant *k*_5_ as proportionality constant. Importantly, continuous sterol uptake and release ensures that the studied cellular system becomes a thermodynamically open system with constant in- and outflow. This allows us to calculate a steady state fraction of sterol in each compartment. Thus, the amount of DHE in each compartment at a steady state 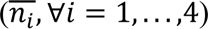 can be analytically derived by setting Eqs 2 and 1b- d to zero and using linear algebra of inhomogeneous equation systems to obtain:

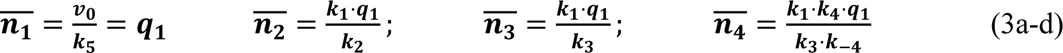

One can see that the steady state ratio in the 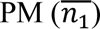 is equal to the ratio of the inflow and release rate constants, and that the steady state sterol amount in all other compartments depends on this ratio as well, i.e., on 𝑞_1_ = 𝑣_0_⁄𝑘_5_, Since these parameters as well as their ratio are not known from the experiments, we seek expressions, that are independent of this ratio and at the same time measurable in experiments. These are the steady state fractions in each compartment 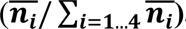, which are given by:

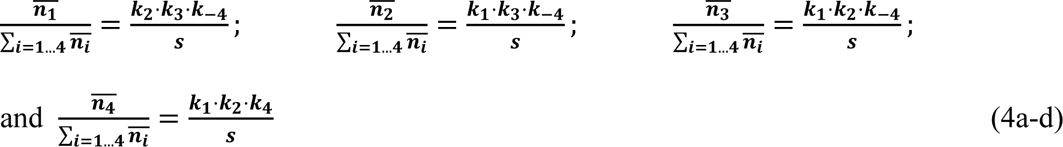

where

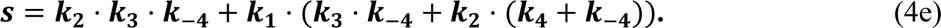

The steady state sterol fraction in the LE/LYSs corresponding to the sum of the fractions of *n*_3_ and *n*_4_ at steady state reads:

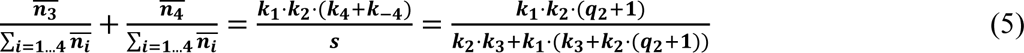

Here, the equilibrium constant between transport to and from ILVs is defined as 𝑞_2_ = 𝑘_4_⁄𝑘_−4_. Similarly, one can define the total intracellular sterol (ignoring minor non-determined fractions in the ER and in mitochondria) by taking the sum of fractional fluorescence of DHE in LEs, ILVs and REs to obtain:

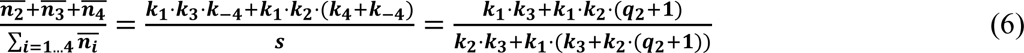

Importantly, the fractions, as determined in Eqs. 4-6 are independent of the influx/efflux ratio *q*_1_. As inferred from Eq. 5 and 6, the sterol fraction in LE/LYSs (including ILVs; compartment 3 and 4) or in REs and LE/LYSs (compartment 2, 3 and 4) drops non-linearly as a function of the rate constant, *k*_-4_. Thus, the higher the transport rate of sterol from ILVs back to the limiting membrane of LE/LYSs, the lower the overall sterol accumulation in cells. One can simplify these expressions by replacing the rate constants *k*_4_ and *k*_-4_ by the equilibrium constant *q*_2_ as in the right-hand side of Eq. 5 and 6. This gives the total intracellular DHE, i.e., the combined amount in REs and LE/LYSs, as shown in Eq. 6, which can be measured from images by intensity thresholding using the ImageJ plugin we presented previously (Berzina et al., 2018). From that, one can compare the calculated sterol fraction in LE/LYSs and REs using the rate constants determined from the kinetic experiments for disease cells (Fig. 2B, square; Fig. 1D and Tab. 1) with the experimentally determined DHE intensities in LE/LYSs + REs from the continuous labeling experiment (Fig. 1B at t=0 from reference (Juhl et al., 2021b)). One finds that the model underestimates the measured sterol fraction by about 20% (i.e. 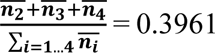 versus intracellular DHE in experiments being equal to 0.495).

This discrepancy could be the result of three processes, which we ignore in our model; first DHE might not only be delivered to the PM in the continuous experiment but instead some DHE could be internalized together with albumin, which we found to be internalized by endocytosis in fibroblasts (Berzina et al., 2018). Second, a minor fraction of sterol delivered to cells as a complex with albumin becomes esterified by ACAT in the ER. By using lipid mass spectrometry of ^13^C-labeled cholesterol complexed with albumin we found that this amounts to about 6% of total labeled cholesterol in control and 0.6% in NPC2-deficient fibroblasts (Berzina et al., 2018). Since, we did not consider such processes in our model, the agreement between experiment and model analysis is remarkable.

A steady state analysis can also be used to predict the behavior of an intracellular transport system for potential parameter changes (Wüstner, 2006b;Wustner, 2019). When plotting the fractional fluorescence of DHE in all three endosomal pools (i.e., the sum of LE/LYSs containing ILVs plus additionally REs; compartment 2, 3 and 4; Eq. 7) at steady state as a function of *q*_2_, we see that cholesterol will accumulate hyperbolically for increasing *q*_2_ (Fig. 2B). Thus, our model predicts that intraluminal trapping of sterol in ILVs due to compromised transport of cholesterol from ILVs back to the limiting endo-lysosomal membrane causes the observed sterol enrichment in endo-lysosomes of NPC2 deficient cells in both the pulse-chase and continuous sterol uptake experiments. The same model analysis can also be used to predict an upper limit for *q*_2_ to restore this transport defect in the presence of NPC2. For that, we ask how much the sterol equilibrium between ILVs and limiting membrane must be shifted to obtain the experimentally determined fraction of DHE in LE/LYSs of control cells. We find that one needs the ratio *q*_2_ to be significantly smaller than one to ensure that the combined sterol fractional in LE/LYS and REs resembles that found in control cells at steady state (Fig. 2B; brown curve).

To test this conclusion, we carried out simulations of the full model with parameters determined for control cells (Tab. 1) and the rate constants *k*_4_ and *k*_-4_ set to two and ten times the value for *k*_4_ found in disease cells, respectively (i.e., *k*_4_ = 0.01686 min^-1^ and *k*_-4_ = 0.0843 min^-1^). This corresponds to a ratio of *q*_2_ = 0.2, and as shown in Fig. 2D, results in good agreement between experiment and simulation. While there is much less sterol in LE/LYSs of control compared to disease cells, there is a comparable amount of sterol in the limiting membrane of LE/LYSs of both cell types (compare Fig. 1D and Fig. 2D, green straight and dashed lines). These results show that the full model can describe the intracellular transport kinetics of DHE in both control, and disease cells very well.

Together, imaging-based kinetic modeling predicts that the affected step in the NPC2 disease state is the intra-endosomal sterol exchange described in our model by the rate constants *k*_4_ and *k*_-4_. Moreover, our analysis suggests that NPC2 increases sterol transport from ILVs back to the limiting late-endosomal membrane (rate constant *k_-_*_4_) more than transport in the opposite direction (i.e., rate constant *k*_4_). As a consequence, cholesterol and other lipids should accumulate in ILVs of NPC2-deficient fibroblasts, but not in fibroblasts from healthy subjects. This would mean, that ILVs should contain more lipid and likely become larger due to the increased lipid content in disease compared to control cells. To test this prediction, high resolution microscopy of lipid content in ILVs versus endo-lysosomal membrane would be needed. Since optical microscopy lacks the resolution to resolve the ultrastructure of LE/LYSs, we employed SXT for this task, as described in the next paragraph.

### 3.2. Soft X-ray tomography of endo-lysosomes reveals lipid enrichment in ILVs of NPC2-deficient cells

SXT is a form of X-ray microscopy gaining increasing attention as an alternative to electron microscopy (EM) for obtaining high resolution ultrastructural information of cellular architecture (McDermott et al., 2009). SXT uses the short wavelength of X-rays of 4.4 nm (corresponding to the K-edge absorption of carbon, 284 eV) to 2.3 nm (the K shell absorption edge of oxygen, 543 eV) to generate absorption contrast in the so-called water window, where carbon-rich material, such as lipid membranes absorb strongly, while water is transparent (McDermott et al., 2009;Schneider et al., 2010). SXT of cryo-frozen cells provides an isotropic resolution of about 40 nm, does not require sectioning and is of higher acquisition speed compared to EM (Schneider et al., 2010). While SXT is based on natural absorption contrast, the technique can be correlated with corresponding fluorescence images, which, however, are mostly diffraction-limited and therefore of much lower resolution (Hagen et al., 2012;Schneider et al., 2012;Duke et al., 2014). We have recently shown that extracellular vesicles, secreted from cells during cholesterol efflux, also contain the cholesterol analogue TopFluor-cholesterol as well as fluorescent NPC2 (Juhl et al., 2021b). For that, 3D reconstructions of X-ray tomograms were aligned with the corresponding cryo fluorescence image of TopFluor-cholesterol. Here, we use the same correlative approach and find by SXT that many intracellular vesicles in human fibroblasts contain ILVs (Fig. 3A). From the corresponding fluorescence images, one finds that the same region is strongly labeled with the fluorescent cholesterol analogue, TopFluor-cholesterol (Fig. 3B-C). Unfortunately, due to the tight packing of vesicles in fibroblasts and the much lower resolution of fluorescence microscopy compared to SXT, we cannot correlate the LE/LYSs between both imaging modalities individually. This limitation might be overcome in the future by combining SXT with super-resolution imaging, such as structured illumination fluorescence microscopy (Kounatidis et al., 2020). Importantly, in NPC2-deficient cells, ILVs as detected by SXT, are much darker and larger than in control cells (Fig. 3D). In contrast to EM, it is more straightforward in SXT to image entire subcellular organelles in 3D. This allows for stepping through the volume of multivesicular LE/LYSs and thereby for assessing the whole population of ILVs. We find by SXT, that endo-lysosomes contain numerous ILVs in both control and disease cells, but those ILVs are darker and larger in NPC2-deficient cells compared to control cells (Fig. 3E and F). The appearance of dark intraluminal membranes in disease cells indicates that they contain more lipids than ILVs in control cells. This can be concluded based on the extent of X-ray absorption, which is linearly proportional to the carbon content (Schneider et al., 2010; Duke et al., 2014; Juhl et al., 2021b; Okolo et al., 2021). Together, these results show that the lack of NPC2 results in strong lipid enrichment, specifically in ILVs in human fibroblasts.

**Figure 3.**
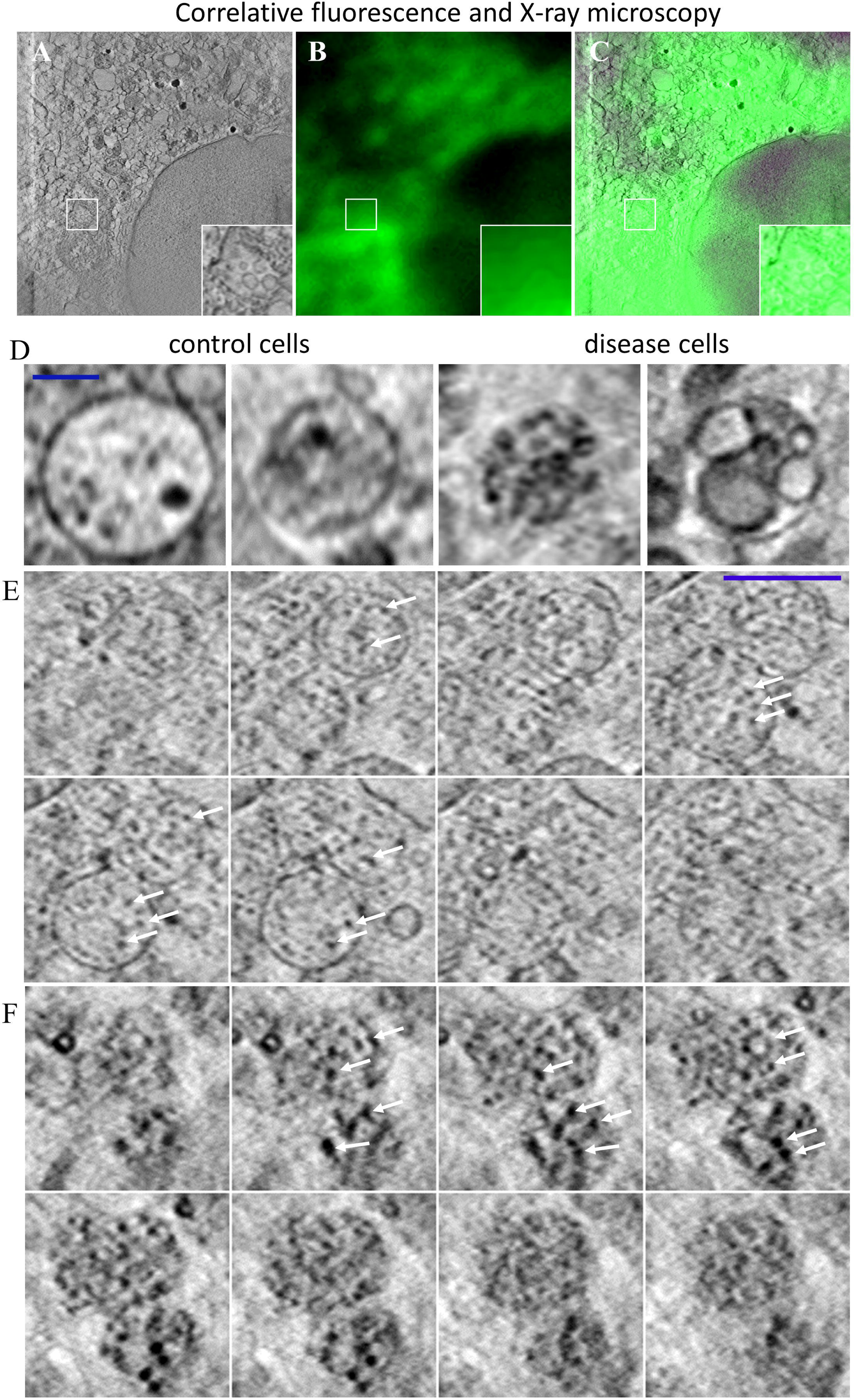
Correlative fluorescence and X-ray microscopy reveal enlarged and lipid-rich intraluminal vesicles in NPC2-deficient fibroblasts. Human fibroblasts were labeled with the cholesterol analogue TopFluor-cholesterol, cryo-frozen and imaged by soft X-ray tomography at the Synchrotron BESSY II as described in Materials and methods. A-C, correlative X-ray and fluorescence imaging of an NPC-deficient fibroblast showing the sum projection of five central image planes from a reconstructed X-ray tomogram (A, inset shows zoomed box with an endo lysosome containing ILVs), the corresponding fluorescence image of TopFluor-cholesterol (B) and the overlay (C). D, two examples of multivesicular LE/LYSs with luminal content in control cells (left two panels) and NPC2-deficient cells (right two panels). E, F, sum projections of three consecutive frames corresponding to a depth of field of 58.8 nm were calculated along a 3D stack reconstructed from an X-ray tomogram of an endo-lysosome in control (E) and disease cells (F). Arrows point to ILVs, which are small and almost transparent in control cells but larger and dark in disease cells due to enrichment of carbon in lipid depositions. See main text for more information.

### 3.3. A lag time model describes sterol efflux from cells via formation of exosomes and ectosomes

The results above show, that the multi-compartment model describes the experimental data on intracellular sterol transport well, and that its predictions are supported by SXT of LE/LYSs in control and disease cells. Our next goal is to determine, whether the model can be used to analyze cholesterol efflux from cells as well. It has been shown in several studies, that when extracellular sterol sources are removed from the medium, NPC disease cells are severely impaired in mobilizing their lysosomal sterol pool compared to control cells (Neufeld et al., 1999;Naureckiene et al., 2000;Rosenbaum et al., 2010). However, it is debated, by which pathway the excess cholesterol can be released from cells, and whether the endo-lysosomal cholesterol destined for efflux must pass through the PM (Rosenbaum et al., 2010;Vacca et al., 2019;Juhl et al., 2021b;Juhl and Wüstner, 2022). We showed recently that human fibroblasts release sterol-rich vesicles of varying sizes during efflux and observed that at least a portion of such vesicles were shed directly from the PM (Juhl et al., 2021b). Such so-called ectosomes have been observed in a variety of cell systems, and they have been associated with cholesterol efflux in several cell types, for example in macrophages and in migrating cancer cells (Ma et al., 2015;He et al., 2018;Juhl and Wüstner, 2022). We found that the abundance of vesicles released from cells correlates with the activity of ABCA1 at the PM, which might work in tandem with NPC2 to export cholesterol from cells (Juhl et al., 2021b). But is this the only pathway for efflux of excess cholesterol from LE/LYSs? Efflux of endo-lysosomal cholesterol from control and NPC disease fibroblasts can be studied using radioactive cholesterol tracers as well as by quantitative fluorescence microscopy in culture medium containing LPDS and without cholesterol source (Yancey et al., 1996;Ko et al., 2003;Pipalia et al., 2006;Lund et al., 2014;Phillips, 2014). Similarly, cells can be loaded with DHE as a cholesterol analogue, and the ratio of fluorescence decrease of DHE in LE/LYSs and in entire cells can be used to determine the sterol fraction in endo-lysosomes over time (Juhl et al., 2021b). Such efflux experiments are typically carried out after steady-state labeling of cells with either deuterated or fluorescent sterols (Haynes et al., 2000;Phillips, 2014;Juhl and Wüstner, 2022). To determine, whether our model, which models sterol efflux from the PM only, can account for measured cholesterol efflux kinetics, we compared first the predicted total cellular sterol at steady state with the measured initial sterol content at the start of an efflux experiment. Our model analysis predicts that the total amount of cholesterol delivered to cells in the continuous uptake experiment is higher in disease compared to control cells. This can be easily seen by summing Eqs. 3a-d to get the total amount of sterol delivered to cells from albumin at steady state:

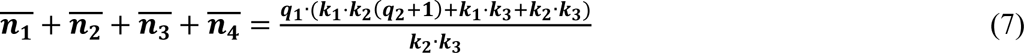

Here, again we have 𝑞_1_ = 𝑣_0_⁄𝑘_5_. By evaluating this expression for the parameter values of control and disease cells, respectively, one finds that the total cell-associated sterol delivered to cells is 4.5fold higher in disease compared to control cells. This is independent from the chosen ratio for the inflow rate and release rate constant from the PM (i.e., independent of *q*_1_), as this parameter drops out when comparing the total sterol delivered to control and NPC2-deficient cells. While being qualitatively correct, this prediction overestimates the difference between control and disease cells, as we found in experiments, that for both, DHE and ^13^C-cholesterol, the sterol amount delivered to NPC2-deficient cells was about 2.5fold that found in control cells (see Fig. 6A in reference (Berzina et al., 2018)). Thus, there must be additional efflux pathways at play apart from cholesterol release from the PM.

Thus, to properly account for the measured kinetics of DHE efflux from endo-lysosomes, we have to consider additional mechanisms apart from sterol release from the PM in our model. Using SXT, we observed not only large ectosomes with a diameter of ≥ 300 nm but also smaller vesicles with diameters of ≤ 50 nm outside of the cell during cholesterol efflux in control and NPC2-deficient fibroblasts (Juhl et al., 2021b). The latter is in the size range of exosomes, which are secreted forms of ILVs released by cells via lysosomal exocytosis, in particular to alleviate lysosomal cholesterol accumulation in NPC disease cells (Strauss et al., 2010). Previous studies using quantitative microscopy of fluorescent cholesterol analogs or of the cholesterol-binding polyene filipin demonstrated that NPC1- and NPC2- deficient fibroblasts can remove excess sterol selectively from LE/LYSs (Pipalia et al., 2006;Rosenbaum et al., 2010;Lund et al., 2014;Juhl et al., 2021b). This would not be expected, if all cholesterol mobilized from LE/LYSs must pass through the PM for efflux. But how can these observations be included in our model?

By measuring the time course of DHE in LE/LYSs, we found that the decay of fractional fluorescence of DHE in endo-lysosomes accelerates over time (Juhl et al., 2021b). We showed that this measured time course of sterol in LE/LYSs can be well described by a stochastic compartment model based on a Weibull survival function in NPC2-deficient cells (Juhl et al., 2021b). This data and the model prediction are replotted in Fig. 4A. The survival function of the Weibull distribution is given by (Macheras and Iliadis, 2006):

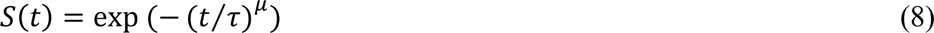

**Figure 4.**
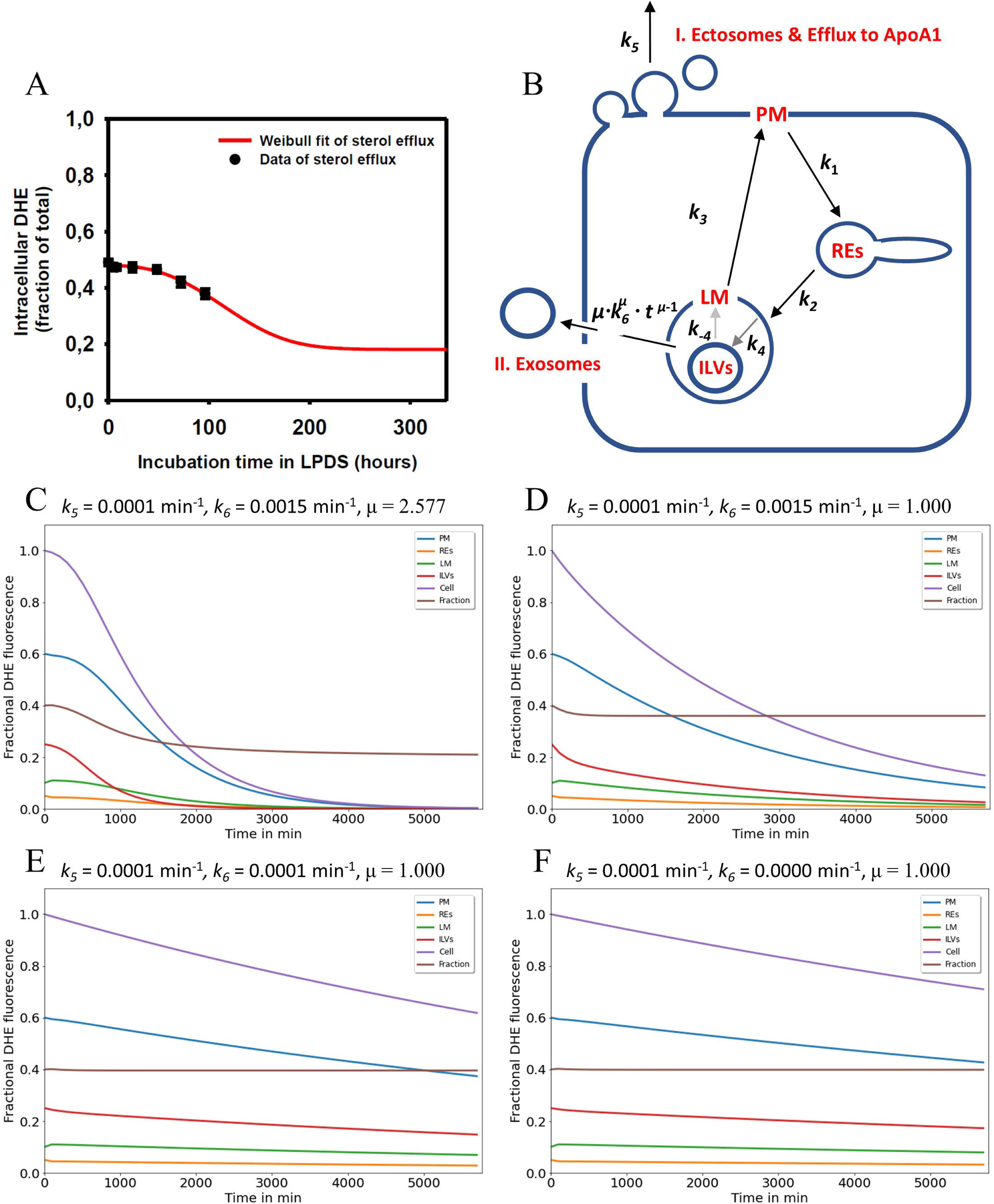
Simulation of cholesterol efflux via two pathways reconciles measured efflux kinetics in NPC2-deficient cells. A, the measured efflux kinetics of DHE from human fibroblasts lacking functional NPC2 is expressed as fractional intracellular intensity (black symbols) and reproduced from Fig. 1 of (Juhl et al., 2021b) with permission. The data is fit to a Weibull survival function (red line) allowing for extrapolation of the efflux kinetics beyond the measured 96h. B, the efflux model is based on the full compartment model and includes additionally sterol efflux from the PM via release of ectosomes and efflux to ApoA1 (I.) as well as sterol efflux directly from LE/LYSs via release of ILVs as exosomes (II.). C-F, simulation of cholesterol efflux from disease cells for different parameter combinations for sterol efflux, i.e., rate constants *k*_5_ and *k*_6_ and the shape parameter μ were varied, while rate constants *k*_1_ to *k*_-4_ were fixed to the values determined for NPC2- deficient cells (see Tab. 1). Simulated time courses are shown for PM (blue lines), REs (orange lines), the limiting membrane (LM) of endo-lysosomes (green lines), the ILVs (red lines) and the total cell (violet lines). In addition, the entire intracellular sterol pool (i.e., sum of REs, LM and ILVs) was calculated and normalized to the total cellular pool (brown lines). This calculated intracellular sterol fraction can be directly compared to the experimentally determined fractional DHE intensity in cells (compare panel A with brown lines in panels C-F). Clearly, the efflux parameter combinations derived for disease cells and shown in panel C fit the data best. See main text for further explanations.

Here, τ is the time constant describing the characteristic time for a sterol molecule to reside (‘survive’) in LE/LYSs before release, and μ is a stretching exponent or shape parameter. For μ > 1 the Weibull survival function models delayed transport processes with an initially delayed but accelerating transport rate, while for μ = 1, a single-exponential decay with constant export rate constant is recovered (Macheras and Iliadis, 2006). The corresponding differential equation describing the rate of change of sterol molecules in LE/LYSs can be found by differentiating the survival function with respect to time, which gives:

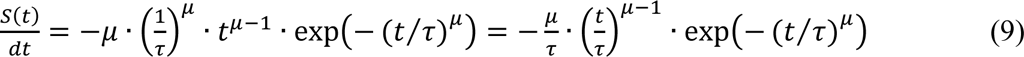

By weighting Eq.8 with the initial fractional fluorescence of DHE in LE/LYSs of NPC2 deficient cells and adding an off set to account for the remaining intensity at steady state, we could estimate efflux parameters for sterol from fibroblasts. We found that this modified Weibull function described sterol efflux very well, providing a time constant for sterol efflux of τ = 135.14h and μ = 2.577 in NPC2 deficient cells (Juhl et al., 2021b). It also allows for extrapolating sterol efflux beyond experimentally accessible time scales, as shown in Fig. 4A (red curve). This model describes the observed phenomenon that lysosomal sterol export is delayed in the beginning of the efflux experiment but accelerates over time. Treating disease cells with purified NPC2 shortened and narrowed the retention time resulting in τ = 86.96h and μ = 1.324 (Juhl et al., 2021b). The Weibull function with µ > 1 is also called a compressed exponential, and a physical motivation for using it in modeling lysosomal sterol efflux of disease cells is the heterogeneous transport of LE/LYSs to the PM before releasing their cargo by exocytosis. We showed previously that endo-lysosomes containing fluorescent NPC2 and DHE undergo a combination of diffusion and active transport in human fibroblasts, justifying the use of this model for analysis of sterol efflux (Lund et al., 2014;Berzina et al., 2018;Juhl et al., 2021b).

To account for this time-delayed and selective efflux of DHE from LE/LYSs, we included two efflux processes into the compartment model, a rate constant describing formation of ectosomes at the PM (*k*_5_), as before (Fig. 4B, pathway I), and a time-dependent rate coefficient *k*(*t*) = 𝜇 ⋅ 𝑘_6_^𝜇^ ⋅ 𝑡^𝜇−1^ to account for the initially delayed sterol release from endo-lysosomes by exocytosis of ILVs as exosomes (Fig. 4B, pathway II). This leads to the following modified equations for the PM (*n*_1_) and the ILVs (n_4_; compare Eqs. 1a, 1d and Eq. 9):

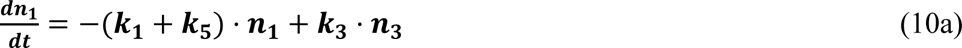

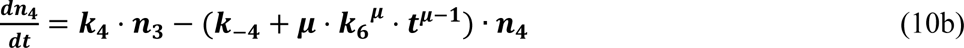

Here, *μ* resembles the shape parameter of the Weibull survival function and *k*_6_ the inverse of the time constant, τ. A numerical simulation of the ODE system consisting of Eqs. 10a, 1b, 1c and 10b for initial sterol fractions in each compartment corresponding to the beginning of the efflux experiment is shown in Fig. 4. Kinetic parameters *k*_1_ to *k*_-4_ were set to the values found for disease cells (see Tab. 1), while the rate constants *k*_5_ and *k*_6_ and the shape parameter μ were varied as shown on top of each panel in Fig. 4C-F. The model does not take homeostatic mechanisms into account which would likely prevent total cellular cholesterol efflux, nor does it account for de novo sterol synthesis, which could replace the effluxed sterol pool. It is instead simply concerned with analysis of the transient kinetics of sterol export from cells, which we determined experimentally in our previous study (Juhl et al., 2021b). Importantly, for μ = 2.577, as previously determined in sterol efflux experiments in NPC2-deficient cells, a characteristic initial delay was found for the sterol amount in the ILVs, the lysosomal membrane and the PM (Fig. 4C, red, green, and blue curve), respectively. The total cellular sterol also decayed after an initial delay phase (Fig. 4C, violet curve). The sum of sterol in ILVs and endo-lysosomal membrane divided by the total cellular sterol gives the sterol fraction in endo-lysosomes (Fig. 4C, brown curve). It shows an initial delay and drops to a steady state value after ca. 200h, corresponding to 1200 min. This is in good agreement with the efflux kinetics we determined previously for disease cells based on parameter fitting of the Weibull function to the efflux kinetics, which could be measured up to 96h (compare Fig. 4A red curve and Fig. 4C, brown curve) (Juhl et al., 2021b). In contrast, this ratio - the sterol fraction in endo-lysosomes - drops exponentially initially and remains constant for longer times, if *µ* = 1.0 for the same remaining parameters (Fig. 4D, brown curve). This is because efflux becomes exponential for both compartments, the PM and LE/LYSs, for *µ* = 1.0, and since *k*_6_ > *k*_5_, the decay from LE/LYSs is slightly faster than sterol efflux from the PM. As a consequence, one finds a slight exponential drop in the ratio of sterol in endo-lysosomes versus total cell. This, however, was not observed in cells (Fig. 4A), showing that the time-delay in lysosomal sterol exocytosis is needed to accurately describe the experimental data. If one sets additionally the rate constant for sterol export from LE/LYSs equal to the one for sterol release from the PM (i.e., *k*_6_ = *k*_5_), the sterol fraction in LE/LYSs becomes a constant rapidly, while the amount of sterol in the PM drops slowly and in an exponential manner (Fig. 4E, brown and blue curve), respectively. If we set the rate constant for sterol efflux from LE/LYSs via exocytosis of ILVs as exosomes to zero (i.e., *k*_6_ = 0), only sterol efflux from the PM remains (Fig. 4F). Importantly, also under those conditions, sterol drops very slowly in all intracellular compartments since they are connected to the PM, from where sterol is released as ectosomes (Fig. 5F, with rate constant *k*_5_=0.0001 min^-1^). However, the sterol fraction in endo-lysosomes stays constant and equal to the initial value at the start of the efflux experiment under these conditions (Fig. 4F, brown curve). These results show that sterol efflux from the PM alone is not sufficient to explain the preferential efflux of sterol from endo-lysosomes, which we observed by measuring the sterol fraction in endo-lysosomes (Fig. 4A, red curve) (Juhl et al., 2021b). It needs an additional mechanism, which removes sterol *selectively* from endo-lysosomes *bypassing* the PM, as otherwise the sterol fraction in endo-lysosomes would be constant despite continuous sterol release from cells. By extending our model to include a second efflux process of cholesterol directly from endo-lysosomes (Fig. 4B, pathway II), we can achieve good agreement between experimentally observed sterol efflux and model predictions. This is a central conclusion of this model analysis. We suggest that this second mechanism is the release of exosomes, i.e., lysosomal exocytosis of ILVs, which has been observed in NPC1-deficient fibroblasts and neuronal cells (Fig. 4B) (Strauss et al., 2010;Demais et al., 2016;Guix et al., 2021;Ilnytska et al., 2021a). This efflux process is well-described by a compressed exponential or Weibull-type export rate. The mechanism underlying this delayed efflux is likely the heterogenous diffusion and transport of LE/LYSs towards the PM followed by lysosomal exocytosis.

**Figure 5.**
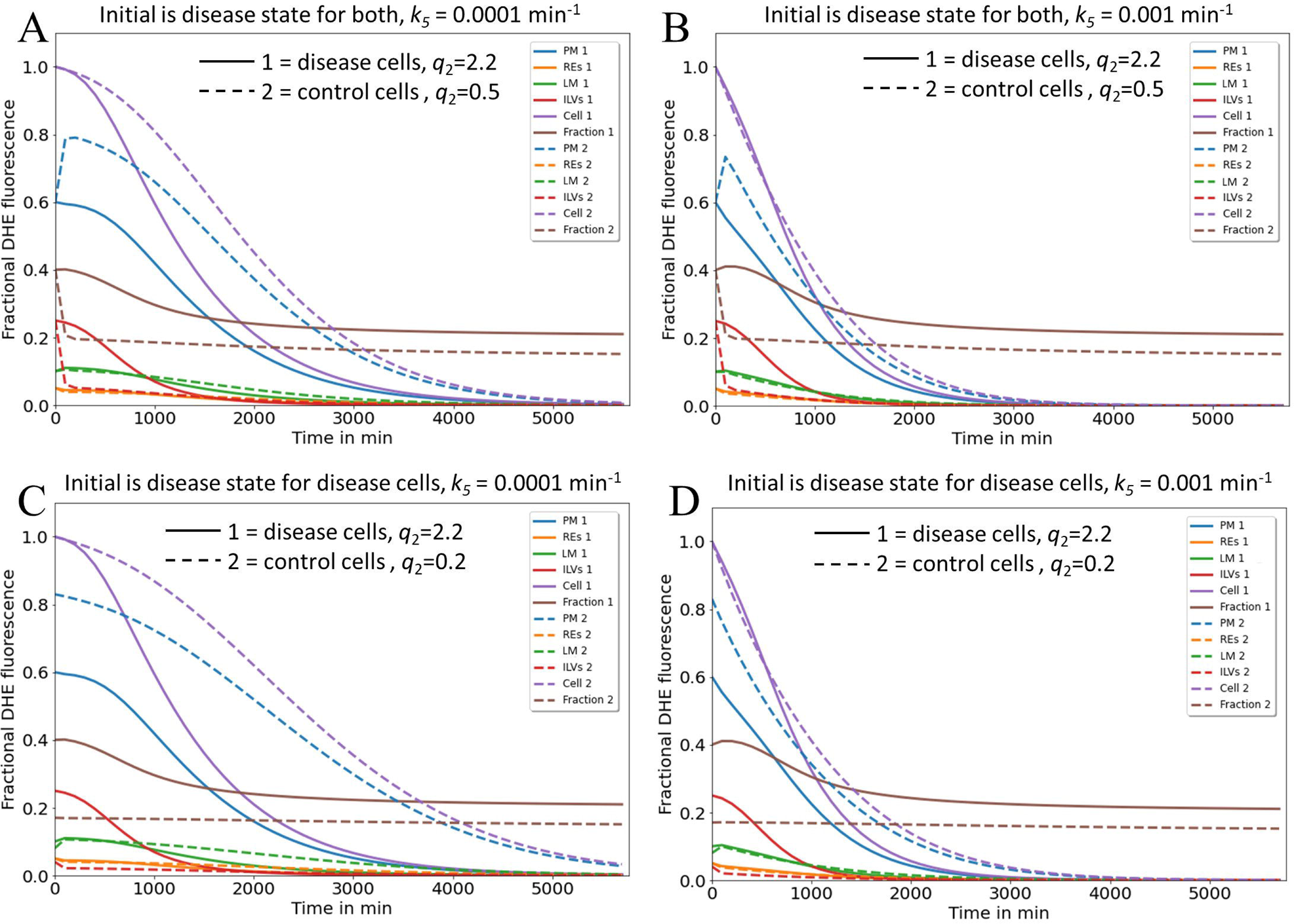
Simulation of cholesterol efflux from control and disease cells predicts reciprocal regulation of the two identified pathways by NPC2. Simulation of cholesterol efflux from disease cells (straight lines) or control cells (dashed lines), either with initial sterol amounts in each compartment as found in disease cells (A, B) or as found in disease and control cells, respectively (C, D). Rate constants *k*_6_ and the shape parameter μ were kept constant at *k*_6_ = 0.0015 min^-1^ and µ = 2.57, while rate constants *k*_1_ to *k*_-4_ were fixed to the values determined for NPC2-deficient cells (straight lines and Tab. 1) or to the values inferred for control cells (dashed lines and Tab. 1). In the latter case, the rate constants *k*_4_ to *k*_-4_ were set to two- and either four- or tenfold the value estimated for *k*_4_ in disease cells, to obtain either *q*_2_=0.5 or 0.2 for control cells. This simulates the more efficient transport of sterol from ILVs back to the limiting membrane of LE/LYSs in the presence of NPC2 and is indicated in the panels. Rate constant *k*_5_, which describes sterol release from the PM, was varied as described on the top of each panel. Simulated time courses are shown for PM (blue lines), REs (orange lines), the limiting membrane (LM) of endo-lysosomes (green lines), the ILVs (red lines) and the total cell (violet lines). Also, the entire intracellular sterol pool (i.e., sum of REs, LM and ILVs) was calculated and normalized to the total cellular pool (brown lines). One finds that total sterol drops faster in disease than in control cells, if the rate constant for sterol efflux from the PM, *k*_5_, is low (C, straight and dashed violet lines). Also, for control and disease cells increasing *k*_5_ ten-fold accelerates the drop in total and PM sterol, but it does not change the intracellular sterol fraction (compare C and D; brown dashed and straight lines). See main text for further explanations.

#### Sterol efflux via release of exosomes and ectosomes could be reciprocally regulated by NPC2 activity

Next, we asked, how these two efflux mechanisms, i.e., sterol release from the PM, either as ectosomes or by lipidation of ApoA1 (pathway I) and sterol efflux by lysosomal exocytosis (pathway II) could be coordinated and how NPC2 might control these processes. For that we simulated the time courses given by the differential equation system consisting of Eqs. 1a, 1d, 10a and 10b with the rate constants determined for disease cells and for control cells, respectively (Fig. 5 straight and dashed lines). Total sterol at the beginning of the efflux simulations was set to one to be able to compare control and disease cells. For rate constants *k*_4_ and *k*_-4_ a two- and fourfold higher value was assumed for control cells than that found in disease cells, which is in line with our analysis of the pulse-chase and continuous uptake experiments (Fig. 2). This will account for faster intraluminal sterol transfer from ILVs to the limiting endo-lysosomal membrane in the presence of NPC2. The initial amount of sterol in each compartment and the *µ*-parameter describing the delayed sterol efflux by release of exosomes corresponded to that found under cholesterol accumulation conditions, i.e., in disease cells.

Under these conditions, we found that the combined sterol fraction in the endo-lysosomal membrane and ILVs normalized to total cellular sterol decay much slower in disease than in control cells (Fig. 5A and B, straight and dashed brown curves). This is in line with the experimentally observed shortening of the sterol retention time in disease cells treated with NPC2 protein (Juhl et al., 2021b). According to our model, the sterol amount in the PM drops in a sigmoid fashion in disease cells but exceeds the initial amount transiently in control cells before eventually decreasing for longer times (Fig. 5A and B, straight and dashed blue curves). Both, the transient increase of sterol in the PM and the rapid sterol decrease in the lysosomal sterol fraction in control cells are a consequence of the faster delivery of the sterol pool from ILVs to the limiting membrane of LE/LYSs in the presence of NPC2. This rapidly replenishes sterol delivered from the lysosomal membrane to the PM in these cells. The faster delivery of sterol to the PM and sterol export from LE/LYSs in control cells cannot be due to faster lysosomal exocytosis, since the parameters for this process are the same for simulations of control and disease cells (i.e., *k*_6_ = 0.01 s^-1^ and µ=2.577). This was chosen by intention to compare the effect of intracellular sterol trafficking on efflux kinetics. In reality, the delay parameter µ could be affected by NPC2, as we observed a narrower retention time of sterol upon treating disease cells with purified NPC2 (Juhl et al., 2021b).

Interestingly, in our simulations total cellular sterol release is more efficient for disease cells, as long as shedding of ectosomes from the PM is slow (Fig. 5A, straight and dashed violet line). We see from that analysis, that secretion of sterol-rich ILVs due to lack of NPC2 mediated intraluminal sterol transfer in disease cells causes a more pronounced net sterol efflux than release of sterol-poor exosomes, found in control cells. Increasing the rate of sterol efflux from the PM, on the other hand, has a proportionally larger effect in control cells, where sterol delivery from ILVs to the limiting membrane of LE/LYSs is more efficient (Fig. 5B, straight and dashed violet line). Still, our model predicts that boosting cholesterol efflux from the PM will also alleviate sterol export in disease cells. This prediction of our model agrees with our recent findings, that upregulation of ABCA1 together with ABCA1 causes a drop in PM sterol but only little change in intracellular cholesterol (see Fig. 8 in (Juhl et al., 2021b)). Only when adding NPC2 protein to the culture medium, intracellular cholesterol could be removed, exactly as predicted by our model.

Supporting the notion of a reciprocal regulation between both efflux pathways are additional simulations, in which we compare the impact of initial conditions in each compartment (Fig. 5C and D). In disease cells, the sterol fraction in LE/LYSs drops in a sigmoid fashion for high initial sterol as found in the absence of NPC2 (Fig. 5C and D, straight brown line). In contrast, in control cells with initial conditions resembling low sterol amounts, no change of the sterol fraction in endo-lysosomes is found (Fig.5C and D, dashed brown line). This is the case despite overall cellular sterol efflux in control cells and for 10-fold varying release rate constants, *k*_5_, from the PM (Fig. 5C and D, dashed violet line).

Our results can be summarized as follows (Fig. 6): in control cells, cholesterol is efficiently transported from ILVs back to the limiting membrane of LE/LYSs, therefore being available for transport to the PM and efflux from cells. Therefore, the first efflux pathway via ectosomes and lipid export to ApoA1 is contributing more to net sterol efflux (Fig. 6A). In disease cells, intraluminal sterol transport in LE/LYSs is hindered and cholesterol accumulates in ILVs due to lack of NPC2. Since ILVs are the precursors for exosomes, more sterol is available for lysosomal exocytosis, i.e., pathway II, in disease cells (Fig. 6B). The latter has indeed been observed in NPC1-deficient cells (Strauss et al., 2010;Demais et al., 2016;Guix et al., 2021). We conclude from this analysis, that sterol efflux by release of exosomes and by shedding from the PM are reciprocally regulated by the intraluminal sterol transfer activity of NPC2 inside endo-lysosomes.

**Figure 6.**
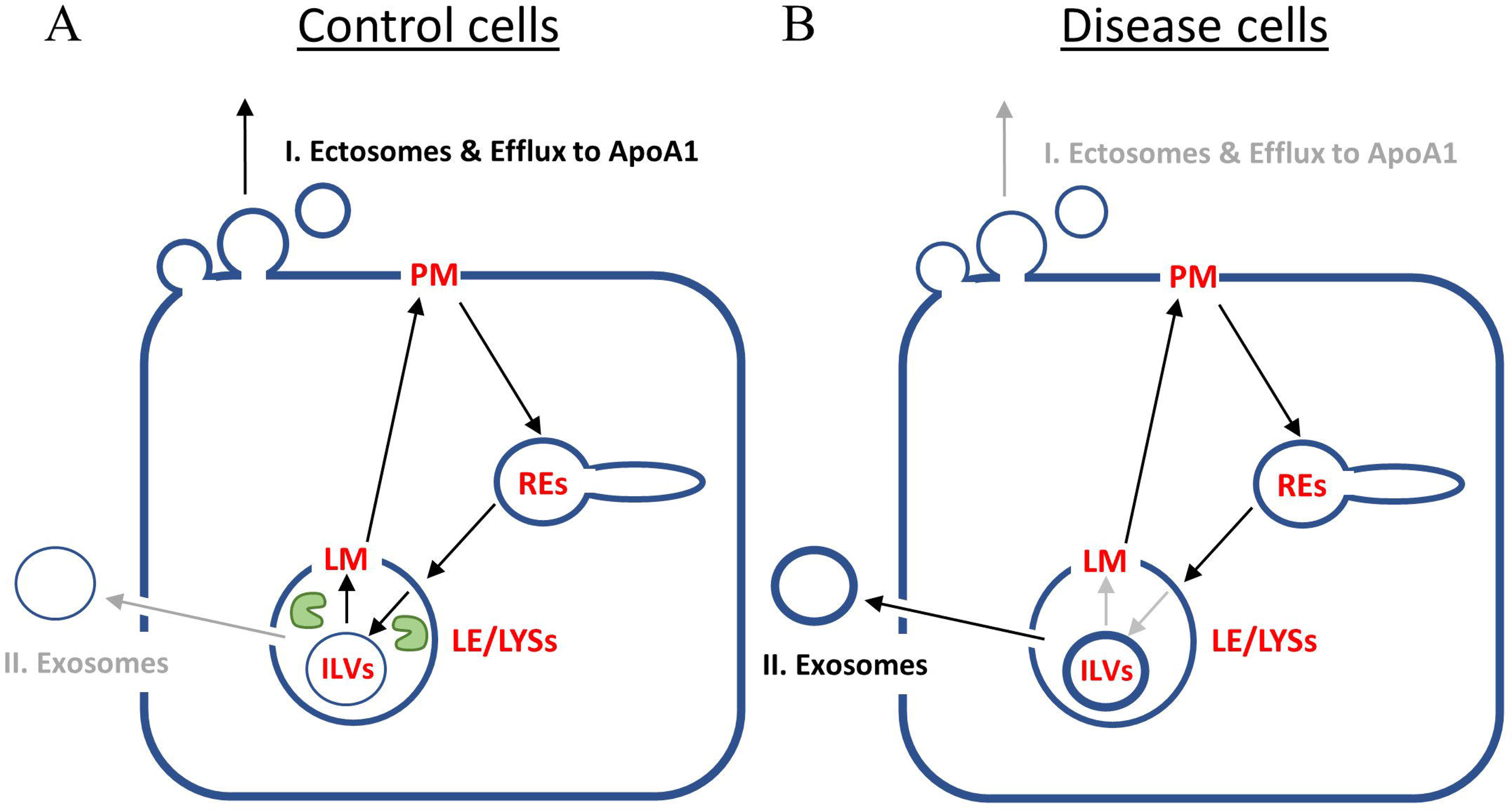
Summary of model predictions on reciprocal regulation of cholesterol efflux pathways by NPC2 activity inside endo-lysosomes. A, in control cells, NPC2 protein inside LE/LYSs (green ‘pac man’) mediates efficient cholesterol transfer from ILVs to the limiting membrane (LM) of endo-lysosomes. This allows for preferred shuttling of sterol back to the PM for efflux via pathway I., i.e., sterol release from the PM as ectosomes and/or by efflux to ApoA1. Therefore, the cholesterol content of ILVs is low (thin blue ring inside LE/LYSs in A), and only little sterol will efflux via pathway II, i.e., lysosomal exocytosis and release of exosomes. B, in disease fibroblasts, the lack of NPC2 results in slow and inefficient cholesterol transfer between ILVs and LM of LE/LYSs. As a consequence, cholesterol build’s up in ILVs (thick blue ring inside LE/LYSs in B). This results in preferred cholesterol efflux via pathway II, i.e., secretion of exosomes, which are nothing but transformed ILVs released from cells. See text for further explanations.

## 4. Discussion

Recent structural and mechanistic studies have provided novel insight into the molecular function of NPC1 and NPC2 in cholesterol export from LE/LYSs. However, the regulation of dynamic sterol shuttling between PM and endo-lysosomes, and the efflux of excess lysosomal cholesterol from particularly NPC2-deficient cells are little understood. Precise quantitative measurements of intracellular sterol transport combined with mathematical modeling of the transport kinetics can provide new insight into these processes. Here, we have developed a mathematical model of sterol trafficking between PM and endo-lysosomes in human fibroblasts. We show that sterol trafficking to LE/LYSs via REs is in line with the available kinetic data on sterol transport in fibroblasts, thereby extending previous modeling attempts for this pathway (Lange et al., 1998). Our model shows that the overall transit time defined as sum of the reciprocal rate constants between PM, REs and endo-lysosomes (i.e. 1/*k*_1_+1/*k*_2_+1/*k*_3_) is about 1.5 h in control cells (Krzyzanski, 2011), supporting earlier suggestions, that LE/LYSs can rapidly respond to changes in PM cholesterol (Lange et al., 1998;Castellano et al., 2017). Our results also support previous findings questioning the general view that initial sterol export from endo-lysosomes is defective in NPC-disease cells (Cruz et al., 2000;Lange et al., 2000). In fact, kinetic modeling suggests that sterol cycling between the limiting membrane of LE/LYSs, and the PM is at least as fast in NPC2-deficient cells as in control fibroblasts (Tab.1 and above). In fact, the transit time for sterol between PM, REs and limiting membrane of LE/LYSs is about 40 min in disease cells, i.e., faster than in control cells, when using the respective mean values of the estimated rate constants, *k*_1_, *k*_2_ and *k*_3_. However, we found that sterol gets trapped inside LE/LYSs leading to its removal from the sterol circuit between PM, REs and limiting membrane of LE/LYSs and instead build up in ILVs inside endo-lysosomes in cells lacking functional NPC2. This results in a second slow phase of sterol accumulation in LE/LYSs of NPC2-deficient cells, not observed in control cells. Using correlative SXT and fluorescence microscopy we were able to confirm enlargement of ILVs with increased lipid content in fibroblasts lacking functional NPC2, thereby supporting our model predictions.

Based on our previous findings (Juhl et al., 2021b), we postulate that efflux of excess endo-lyosomal cholesterol from fibroblasts can take place via two pathways, I) efflux from the PM, at least partly involving ectosome release and II) secretion of exosomes in a form of lysosomal exocytosis. We model the latter pathway via a Weibull-type differential equation, which allows us to account for the delayed decrease in the intracellular sterol fraction upon removal of sterol sources from the culture medium (Juhl et al., 2021b). Delays in cholesterol efflux have also been included in a previous study in macrophages, but the authors did this by postulating additional transit compartments (Gaus et al., 2001). This could also be done in our system. In fact, reasonable fits were also found for modeling lysosomal sterol export using an Erlang-type distribution in the efflux experiments (not shown). However, this requires postulating additional compartments, for which no experimental evidence is currently available.

Our modeling results suggest that both efflux pathways could be reciprocally regulated by the sterol transfer activity of NPC2 between ILVs and the limiting membrane of endo-lysosomes: in cells lacking NPC2, the diminished transfer capacity between ILVs and the limiting membrane causes cholesterol build up in ILVs, which the affected cells clear via preferred sterol efflux by exosome secretion. In contrast, in control cells, cholesterol transfer from ILVs to the membrane of LE/LYSs is catalyzed by NPC2 resulting in efficient sterol transfer to the PM, such that the first pathway of sterol efflux should be used preferentially (Fig. 6). These predictions are supported by experimental findings in NPC1 deficient neurons and glia cells, where not only more and larger ILVs were found but also more exosomes were secreted than in control cells (Strauss et al., 2010;Guix et al., 2021;Ilnytska et al., 2021b). Supplementation of NPC1-deficient fibroblasts with phosphatidylglycerol, the precursor of LBPA and other lipid species in ILVs, caused an increased release of cholesterol-rich exosomes and a down-regulation of ABCA1 (Ilnytska et al., 2021a). The latter is expected if reciprocal regulation of sterol efflux from the PM via an ectosome/ABCA1 pathway and lysosomal exocytosis of exosomes is at play in cellular cholesterol efflux. LBPA, which is formed from supplemented PG, is abundant in LE/LYSs, where it can contribute up to 15-20 mol% of total phospholipids. It is likely, that LBPA location and abundance inside LE/LYSs plays a key role in this reciprocal regulation; it is not only a binding partner for NPC2, enhancing interbilayer sterol transfer by NPC2 but also an activator of acid sphingomyelinase, which converts sphingomyelin into ceramide (Oninla et al., 2014). Ceramide, on the other hand, is a lipid with high negative spontaneous curvature due to its low head-group to acyl chain ratio. It is directly involved in intraluminal budding of vesicles and thereby formation of ILVs inside endo-lysosomes (Trajkovic et al., 2008). Interestingly, production of ceramide from degradation of sphingomyelin by acid sphingomyelinase increases NPC2 mediated sterol transfer between liposomes severalfold (Abdul-Hammed et al., 2010;Oninla et al., 2014). Incubating NPC1-deficient fibroblasts with purified acid sphingomyelinase ameliorates cholesterol clearance and trafficking capacity in these cells (Devlin et al., 2010).

Based on these findings it is reasonable to speculate that the gradient of these lipids, LBPA and ceramide, inside LE/LYSs could also facilitate directional sterol transfer by NPC2 between ILVs and the limiting membrane of endo-lysosomes. This, in turn, would contribute to the reciprocal regulation of the two sterol efflux pathways. Coupling of sterol transfer to local metabolism and turnover of a second lipid has been proposed as a mechanism for selective sterol enrichment in the Golgi by oxysterol binding proteins, which shuttle sterol and phosphatidyl-inositol-4-phosphate directionally between Golgi and endoplasmic reticulum (ER) (Mesmin and Antonny, 2016). In a similar way, NPC2 in concert with LBPA and ceramide might control the sterol distribution between endo-lysosomal membrane and ILVs and thereby influence, which cholesterol efflux pathways from LE/LYSs is primarily used. Future experiments should test the model proposed here, for example by studying intracellular trafficking and lysosomal export of cholesterol analogues in disease fibroblasts treated with PG, LBPA or acid sphingomyelinase. Similarly, sterol transport kinetics could be analyzed with our model in cells treated with mutated NPC2, in which the interaction with LBPA and with sterol transporters such as NPC1 or LAMP2 has been removed (Deffieu and Pfeffer, 2011;Li and Pfeffer, 2016;McCauliff et al., 2019). Finally, it would be very interesting to explore the link between NPC2 and lysosomal exocytosis further, as this process is known to be regulated by calcium-mediated signaling via the endo-lysosomal ion channel mucolipin-1 as well as the transcription factor TFEB, a master regulator of lysosome biogenesis (Medina et al., 2011;Shen et al., 2012;Vacca et al., 2019). In conclusion, our study shows that quantitative modeling of intracellular sterol transport and efflux combined with high-resolution multimodal imaging plays an important role in elucidating mechanisms of lipid transfer and in testing hypotheses of cellular lipid homeostasis.

## 5. Author Contributions

DW designed the study, developed the mathematical model, fit it to the data and wrote the first draft of the manuscript. ADJ prepared and labeled cells and carried out SXT measurements with help of SW, JM, and GS. JME together with ADJ and DW analyzed the X-ray image data. All authors red and approved the final version of the manuscript.

## Data Availability Statement

The datasets generated and analyzed for this study can be found in the [NAME OF REPOSITORY]

## Supplemental figure legends

**Figure S1. Alternative models for the pulse-chase kinetic data of DHE transport**

In the parallel model shown in A, DHE traffics bi-directionally in two branches originating at the plasma membrane (PM); 1) to late endosomes and lysosomes (LE/LYSs; left branch in A) or to the recycling endosomes (REs; right branch in A). In the sequential model shown in B, DHE traffics sequentially from the PM to the REs and from there to the LE/LYSs. Both models were solved numerically in SAAM software and fitted to the experimental time courses, described in Fig. 3D, for control cells (C) and for NPC2 disease cells (D). The symbols show the experimental data of the DHE fraction in the PM (black circles, control cells; black triangles, disease cells), in the REs (green circles, control cells; green triangles, disease cells) and in the LE/LYSs (red circles, control cells; red triangles, disease cells). Black and brown straight lines show the fit of the parallel and sequential model, respectively, to the DHE data in the PM. Green and spinach green straight lines show the fit of the parallel and sequential model, respectively, to the DHE data in the REs. Red and orange straight lines show the fit of the parallel and sequential model, respectively, to the DHE data in the LE/LYSs. Dotted lines in D indicate a fit with the parallel model but with rate constants for the second branch between PM and REs constraint to the values found in control cells (compare green line in C). This constraint did not affect the fit of the parallel model to the time course of DHE transport from the PM to the LE/LYSs, showing the independence of both branches in this model. Overall, the fit quality of both models, as judged by the Chi-square value and Akaike criterion, was much worse compared to that for the model shown in the main text.

