## Supplementary figures and images for "Kinetic modelling of sterol transport between plasma membrane and endo-lysosomes based on quantitative fluorescence and X-ray imaging data"

### Supplemental Figure 1

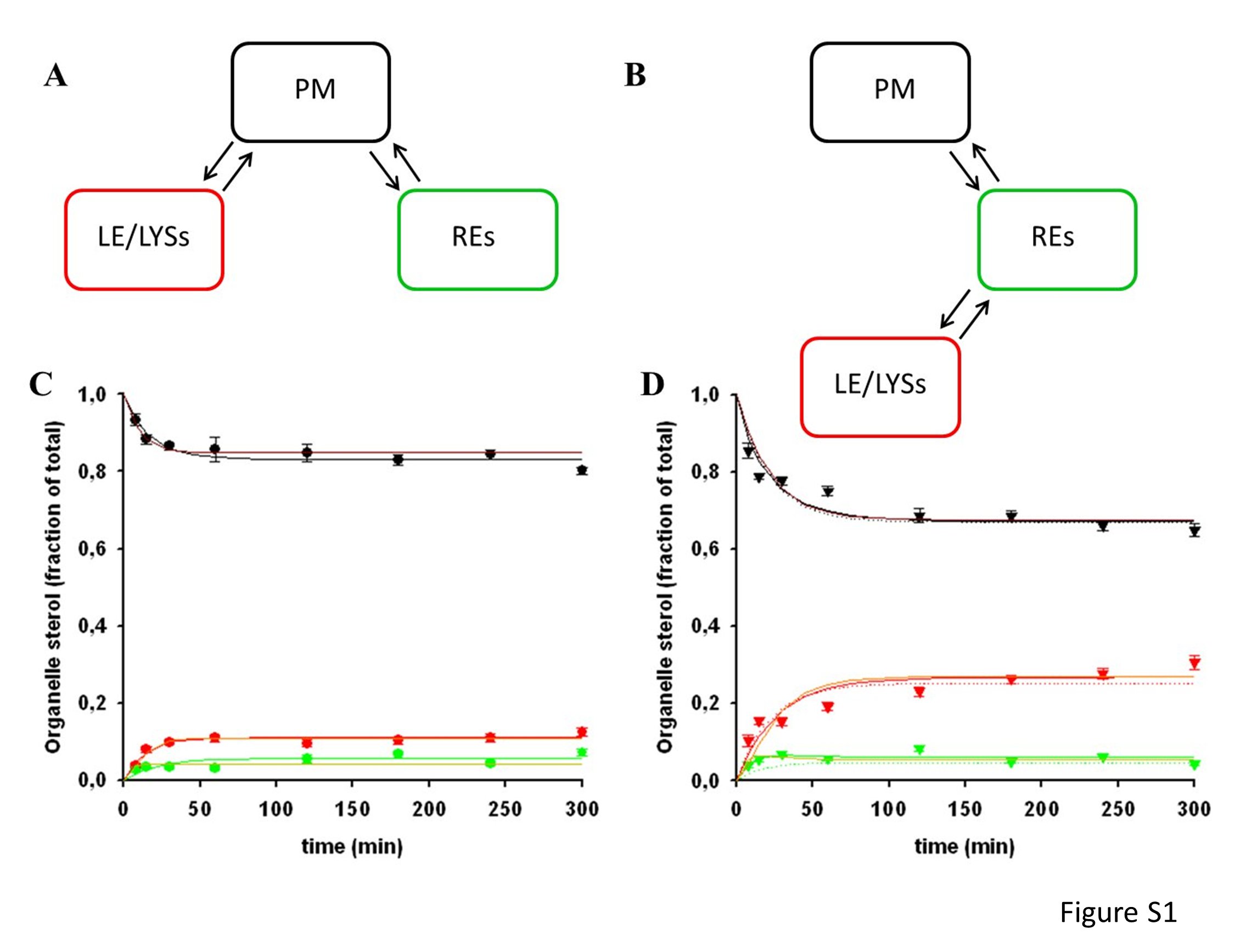
